# Old data and friends improve with age: Advancements with the updated tools of GeneNetwork

**DOI:** 10.1101/2021.05.24.445383

**Authors:** Alisha Chunduri, David G. Ashbrook

**Affiliations:** Department of Biotechnology, Chaitanya Bharathi Institute of Technology, Hyderabad 500075, India; Department of Genetics, Genomics and Informatics, University of Tennessee Health Science Center, Memphis, TN 38163, USA

## Abstract

Understanding gene-by-environment interactions is important across biology, particularly behaviour. Families of isogenic strains are excellently placed, as the same genome can be tested in multiple environments. The BXD’s recent expansion to 140 strains makes them the largest family of murine isogenic genomes, and therefore give great power to detect QTL. Indefinite reproducible genometypes can be leveraged; old data can be reanalysed with emerging tools to produce novel biological insights.

To highlight the importance of reanalyses, we obtained drug- and behavioural-phenotypes from Philip *et al.* 2010, and reanalysed their data with new genotypes from sequencing, and new models (GEMMA and R/qtl2). We discover QTL on chromosomes 3, 5, 9, 11, and 14, not found in the original study.

We narrowed down the candidate genes based on their ability to alter gene expression and/or protein function, using *cis*-eQTL analysis, and variants predicted to be deleterious. Co-expression analysis (‘gene friends’) and human PheWAS were used to further narrow candidates.

Prominent candidate genes include: *Slitrk6* in a Chr 14 QTL for locomotion in the center of an open field, we show to be part of a coexpression network involved in voluntary movement, and association with neuropsychiatric phenotypes in PheWAS; and *Cdk14*, one of only 3 genes in a Chr 5 QTL for handling induced convulsions after ethanol treatment, that is regulated by the anticonvulsant drug valproic acid.

By using families of isogenic strains, we can reuse and reanalyse data to discover novel and highly plausible candidate genes involved in response to the environment.

## Introduction

Two of the biggest problems in analyses of biomedical data are irretrievability and irreplicability. Biomedical data is often lost as soon as it is published, locked within a forgotten harddrive, or siloed in a little-used format on a lab's website. There are many efforts to make data publicly retrievable, such as the FAIR Principles [1], and these are allowing the combined analysis of many datasets, and reanalysis using new tools. There is still the problem of irreducible datasets: for example if a sample from a particular outbred cohort is found to be an outlier during data analysis, there is no way to go back to that genometype and remeasure the phenotype. Nor can new phenotypes be measured in the same individuals in the same environments at a later date as new tools emerge. Families of isogenic strains solve this problem, giving us reproducible genometypes that can be sampled many times, under many environmental conditions: so called experimental precision medicine [2]. This means that a genometype sampled 30 years ago in a different country, can be replicated now, in any lab, with any environmental variable of interest, using any technique. The GeneNetwork.org website allows this combination of FAIR data and reproducible genomes, meaning that we can now go back to previous datasets, and reanalyze them with new data and new tools. Every new dataset adds exponentially to the number of possible connections. In this paper, we will reanalyze drug and addiction related data from over a decade ago using new genotypes for the BXD family and new tools, showing that we can identify new quantitative trait loci (QTLs) and highly plausible candidate genes.

Quantitative trait locus (QTL) mapping has been carried out in numerous species to associate regions of the genome to phenotypes, even before the structure of the genome was well understood (e.g. [3]). Rodents, especially mice, have been the species most prominently used for biomedically relevant traits. Amongst these, the BXD family of recombinant inbred (RI) mouse strains, have been extensively used for almost 50 years in fields such as neuropharmacology [4,5,6,7,8,9,10], immunology [11,12,13,14,15], behavior [16,17,18,19,20,21,22,23], aging [24,25,26,27,28], neurodegeneration [29,30,31,32,33], and gut microbiome-host interactions [34].

The development of the BXD panel was started by Benjamin A. Taylor by inbreeding the progeny of female C57BL/6J and male DBA/2J strains, for the purpose of mapping Mendelian traits [35]. This led to the original 32 BXD strains, which now carry the suffix ‘TyJ’ (**T**a**y**lor to **J**ackson Laboratory). To increase the power and precision of QTL mapping, the number of strains has been expanded [36], including through advanced intercross by [37], and now to a total of 140 extant strains [2], making this resource the largest family of murine isogenic strains. Phenotypes in the BXD have been measured under many conditions, allowing identification of gene-by-environment interactions. Understanding these interactions can potentially help in the discovery of complex therapeutic solutions, and are a vital part of the development of precision medicine.

GeneNetwork.org is a tool for quantitative genetics that started in 2001 as WebQTL [38]. It evolved from analyses of forward genetics in the BXD mouse family, to phenome-wide association studies and reverse genetics in a variety of species. Although GeneNetwork contains data for many species and populations, it most prominently contains data for the BXD family. Over 10,000 ‘classical’ phenotypes, measured under a variety of environmental conditions, and over 100 ‘omics datasets are available on GeneNetwork for the BXD family. GeneNetwork and BXD RI population are therefore a powerful tool for systems genetics and experimental precision medicine.

The great advantage of inbred lines, with stable genome-types that can be resampled, is that data can be reused and reanalyzed over time, as tools improve. From the very start of the genome sequencing revolution, when loci were first mapped to causative genes, new tools and a greater understanding of the genome have allowed us to go back to old data and gain new insight.

In this study, we will demonstrate how new biological insight into drugs of abuse can be gained by reanalyzing data in the BXD family, using improved genotypes from sequencing, and new mapping methods (linear mixed models). Using this method, we have discovered new QTLs and candidate genes for behavioral phenotypes associated with the predisposition of drug- and behavior-related traits obtained from Philip et al. 2010 [39].

## Methods

### Phenotype data

The traits used for analysis in this study were acquired by Philip, and the team and published in 2010 [39]. All data from this publication are freely available on GeneNetwork.org, and were obtained from the BXD published phenotypes. Philip’s study aimed to determine the influence of genes in the response to environment and plausibility of similar interaction with the drug-related attributes including response to and, withdrawal from cocaine, 3,4-methylenedioxymethamphetamine, morphine, and ethanol and their correlation to traits including anxiety, locomotion, stress sensitivity, and pain sensitivity. Complex phenotyping batteries consisting of diverse behavioral assays were employed on the RI strains and multi-variate analyses were performed using GeneNetwork.org. An interplay between environmental factors, drug-induced neural changes, and genetic factors underlie the predisposition of an individual to addiction. In our study, a total of 762 traits were analyzed [Supplementary table 1] using new genotypes and the GEMMA mapping software, to identify novel candidate genes and gene-by-treatment interactions. We did not include morphine related traits, as these are being actively studied by others. Of the then extant population of 79 strains [7], Philip’s study used approximately 70 strains to measure the traits.

### New genotypes from sequencing

A total of 152 BXD strains have now been sequenced using linked-read technologies, and new genotypes for all 152 BXD strains have been produced from this (Personal communications). Variants were chosen to define the start and end of each haplotype block, and variant positions from the previously published genotypes were kept to allow maximum back compatibility with previous publications.

### GEMMA, kinship within the BXD and QTL mapping

The BXD family has been produced in several ‘epochs’ across 40 years, using both standard F2 recombinant inbred methods, and advanced intercross recombinant inbred methods [2]. This has led to both expected and unexpected kinship between BXD strains. This kinship between strains can lead to bias, as it breaks the expectations of previously used methods, such as the Haley-Knott mapping algorithm that was used by Philip’s study. Updated linear mixed models including Genome-wide Efficient Mixed Model Association (GEMMA) and R/qtl2 accessible in the GeneNetwork have been used for this study, as they allow correction for kinship, as well as other cofactors if needed.

Analysis of the 762 traits (taken from Philip et al. study) was carried out using the GEMMA mapping tool with the genotypes from sequencing, a minor allele frequency (MAF) of 0.05 and utilizing the Leave One Chromosome Out (LOCO) method. This computation provides a −log(p) value between each marker and the phenotype. We used a −log(p) > 4, as significant. However, since permutations of the GEMMA algorithm are not currently available in GeneNetwork, we confirmed the significance of these QTL using the linear mixed model tool within r/QTL2 [40], with 5000 permutations of the data.

### Identification of novel QTL

Two methods were used to identify significant QTL.

Firstly, traits with an adjusted p < 0.05 using permutation in r/QTL2 (described above), as these are significant after empirical correction.

The second method we used was to take advantage of independent traits which share QTL at the same location with suggestive p-values (p < 0.63). This p < 0.63 equates to one false positive per genome scan. However, the likelihood of any particular chromosome having a QTL on it is approximately 1 in 20 (i.e. p < 0.05) due to there being 20 chromosomes in mice. The likelihood of two independent traits sharing the same QTL location by chance is therefore much lower than p < 0.05. Traits were called as independent if they were carried out in separate groups of animals (e.g. males and females) or if the traits were measured at independent timepoints (e.g. at 10 minutes after treatment and 60 minutes after treatment).

### QTL confidence intervals

A 1.5 LOD or 1.5 −log(p) drop was used to determine the QTL confidence interval for each statistically significant trait (in our particular case of a two-parent population LOD and −log(p) are approximately equal). Therefore, for each of the QTL above [Supplementary table 2] we were able to generate a list of genes within this confidence interval. Genes were called within our QTL interval using the genenetwork QTL mapping tool - this provides both protein coding genes, non-coding genes, and predicted gene models.

### *Cis*-eQTL mapping

A *cis*-eQTL indicate that a variant within or very close to a gene influences its expression. Genes with cis-eQTLs are high priority candidates, as it provides a potential causal pathway between the gene variant and the phenotype of interest (i.e. the variant alters gene expression, and the expression of that gene alters the phenotype). Therefore, if a gene within a QTL interval is cis-regulated, we catagorize it as a high priority candidate.

For each QTL, we identified which, if any, genes within the QTL interval also had a *cis*-eQTL, and in which tissues an eQTL was seen (using transcriptome data from GeneNetwork). Using this same data, we also identified correlations between expression of each of these genes and the phenotype of interest.

### ‘Gene Friends’, or coexpression analysis

Genes with a *cis*-eQTL in at least one tissue were further considered for coexpression analysis. The top 10,000 correlations were generated in the tissue with the highest correlation between gene expression and the phenotype of interest. Gene-gene correlations with Sample p(r) < 0.05 were taken into Webgestalt to perform an over-representation analysis [41, 42, 43, 44]. This results in identification of significantly enriched annotations or pathways, in the genes which co-express with our gene of interest. This can often suggest pathways or networks that the gene is involved in, even if the gene itself has not yet been annotated as part of that network.

### Gene variant analysis

Deep, linked-read sequencing of the 152 members of the BXD family has been carried out using Chromium 10X sequencing (https://www.10xgenomics.com/products/linked-reads), resulting in 5,390,695 SNPs and 733,236 indels which are high confidence, and segregate in the population (i.e. have a minor allele frequency greater than 0.2). These 6 million variants are potential causes of QTLs detected in the BXD family.

To identify potential effects of these variants, we used the Variant Effect Predictor (VEP) website (http://ensembl.org/Tools/VEP and [45]). All variants within our QTL intervals were extracted from the variant vcf file, and uploaded to VEP. Potentially deleterious variants or variants which impact protein function were identified using the ‘Consequence’, ‘IMPACT’, ‘SIFT’ [46,47] and ‘BLOSUM62’ [48] annotations.

### PheWAS

Phenome-wide association studies (PheWAS) take a genomic region of interest, and find associations between that region and phenotypes measured in GWAS datasets. We used human PheWAS data for all of the candidate genes in our QTLs to detect the genes with relevant human phenotype associations (i.e. behavioral and neurological phenotypes). A relevant association implies confidence in a candidate gene, and suggests cross-species translatability of the finding. We used online PheWAS tools, GWASatlas (https://atlas.ctglab.nl/PheWAS, [49]) and PheWeb (http://pheweb.sph.umich.edu/) for this study.

## Results

### Identification of QTLs

We first sought to identify novel genetic loci linked to the phenotypes from Philip et al., 2010 [39], that were not found in the original study. Comparing QTL mapping using Haley-Knot (H-K; as used previously) and GEMMA, there are 426 traits which had a maximum LRS < 17 with H-K (i.e. non-significant), that now have a maximum −log(p) > 4. These new QTL are therefore of interest.

To confirm these, we performed linear mixed model (LMM) QTL mapping in R/qtl2, with permutations. This produced 61 traits which are significant compared to the empirical significance threshold generated by permutations [Supplementary table 3].

Two methods were used to identify QTL of interest. First, we had the group of 61 traits that were significant by permutations. The second method we used was to take advantage of independent traits which share QTL at the same location with suggestive p-values (p < 0.63). Traits were called as independent if they were carried out in separate groups of animals (e.g. males and females) or if the traits were measured at independent timepoints (e.g. at 10 minutes after treatment and 60 minutes after treatment). We identified 25 QTL for 267 traits [Supplementary table 4].

### Novel QTL

For each of the QTL identified above, we determined if they were reported in Philip et al’s original study [39], or if related phenotypes have been reported in the MGI database [50].

Several locomotion related QTL map to Chr1:37.671-78.94 Mb, that were not detected in the Philip *et al.* study. Previously detected relevant phenotypes associated with this region include the loss of righting induced by ethanol 1 QTL [51] and a vertical clinging QTL [52].

We report a novel QTL on Chr3 (51.723-56.473 Mb) for vertical activity, and on Chr 4 (105.245-114.11 Mb) for locomotion in response to cocaine. Previous studies show a QTL for anxiety in this region of Chr 4 [53]. We also report novel QTLs on Chr 5 for handling induced convulsions as an ethanol reponse (4.468-5.172 Mb) and locomotion in response to cocaine (99.801-101.331 Mb). Finally, there was a novel QTL for locomotion in response to cocaine on Chr 11 (46.361-50.383 Mb) [Supplementary table 4]. Philip’s study shows the presence of multiple significant QTL on Chr 13 that include locomotor (Injection stress-induced activation), morphine withdrawal measures such as jumps, defecation and urination, response to sensitivity and anxiety related to acute stress.

### Candidate causal genes within novel QTL

We concentrated on a subset of six novel QTL which contained less than 100 genes. These QTL are more amenable to finding plausible candidate genes using bioinformatic methods. We reduce the likelihood of finding false positives, and these large QTL are more likely to be due to two or more variants in different genes both contributing to the phenotype. The advantage of families of isogenic strains, like the BXD, is that more strains could be phenotyped, reducing the size of these QTL regions, and allowing greater precision. We leave these large QTLs to future studies.

The ‘small’ QTL we use as examples here were: Chr3:51.723-56.473 Mb, Chr 5:4.468-5.172 Mb, Chr5:99.801-101.331 Mb, Chr9:45.671-48.081 Mb, Chr11:62.923-65.082 Mb and Chr 14:109.994-114.751 Mb [Supplementary table 4]

We used several tools to narrow down potential candidate genes within these QTL. Variants can change phenotype in two main ways: they can either change gene expression, or can change protein function.

To look for variants altering gene expression, we first looked for genes within our QTL regions which has a local or *cis*-eQTL. *Cis*-eQTL demonstrate that there are variants in or close to a gene that cause changes in that gene’s expression. This is useful, since it clearly shows that a variant in the eQTL region has a regulatory effect. Therefore, genes with a *cis*-eQTL are interesting candidate genes.

The next step is to see if expression of these genes correlates with the phenotype(s) of interest. This would suggest a chain of causality: A variant within a gene causes a change in its expression, and the expression of that gene correlates with expression of a trait of interest. To do this, we created a correlation matrix between all genes within a QTL with a *cis*-eQTL in any brain tissue [Supplementary table 6] and the phenotypes that contributed to the QTL [Supplementary table 6]. Any gene with a *cis*-eQTL and a significantly correlated expression was considered a good candidate. If the gene only had a *cis*-eQTL and correlation in a single brain region, then it suggested that this brain region might also be of interest for the phenotype (adding another link to this chain).

The QTL region for vertical activity (Chr 3 51.723-56.473 Mb), has 60 genes among which six genes show presence of *cis*-eQTLs [Supplementary table 6]. No relevant functional annotations (Gene Ontology) have been reported yet. *Dclk1* (location of cis-eQTL: Chr3 55.52 Mb) variants were previously reported to be associated across Schizophrenia and Attention Deficit Hyperactivity Disorder [54]. The same gene has been described as a candidate gene for inflammatory nociception [55]. *Trpc4* (location of cis-eQTL: Chr 3 54.266176 Mb) may be involved in the regulation of anxiety-related behaviors [56].

The QTL region for handling induced convulsions (ethanol response; Chr 5 4.468-5.172 Mb), two genes (*Fzd1* and *Cdk14*) with *cis*-eQTLs among the three present in this region. *Fzd1* (location of cis-eQTL: Chr 5 4.753 Mb) receptor regulates adult hippocampal neurogenesis [57].

The locomotion in response to cocaine QTL (Chr 5 99.801-101.331 Mb) has ten genes with *cis*-eQTLs. QTL Analysis of *Enoph1* (location of *cis*-eQTL: Chr 5 100.062 Mb) in mice indicates that it plays a role in stress reactivity [58]. Variants of *Coq2* (location of *cis*-eQTL: Chr 5 100.654 Mb) contribute to neurodegenerative disorders like Parkinson’s disease [59]. Relevant annotations for other genes with *cis*-eQTLs have not been reported yet by other studies.

The mechanical nociception QTL (Chr 9 45.671-48.081 Mb), incudes five genes with *cis*-eQTLs. *Sik3* (location of *cis*-eQTL: Chr 9 46.222 Mb) is involved in regulating NREM sleep behavior in mice [60]. *Cadm1* (location of *cis*-eQTL: Chr 9 47.550 Mb) knockout mice show increased anxiety, impaired social and emotional behaviors and disrupted motor coordination [61].

Analysis of the locomotion in response to cocaine QTL (Chr 11 62.923-65.082 Mb) revealed five genes with *cis*-eQTLs. *Arhgap44* (location of *cis*-eQTL: Chr 11 65.005456 Mb) has phenotype associations related to abnormal motor learning, abnormal response to novel object, increased grooming behavior and hypoactivity [62]. The brain regions with highest correlation have been added to the supplementary information [Supplementary table 6].

The locomotion in the center QTL (Chr 14 109.994-114.751 Mb), has a single gene with a *cis*-eQTL, *Slitrk6* (location of *cis*-eQTL: Chr 14 109.231826 Mb). Knockout of this gene has been associated to impaired locomotory behavior and altered responses to a novel environment making this gene a strong candidate [63].

### Coexpression networks or ‘gene-friends’

Genes that are coexpressed are often parts of the same pathways or networks, contributing to similar phenotypes. These so-called ‘gene-friends’ [64,65] can provide insight into the function of an unannotated gene and can be implied from the annotated functions of the genes it coexpresses with. As new omics data are being generated for the BXD all the time (now including methylation, proteomic and metabolic datasets), new gene-friends can be found.

For each of the genes within our QTLs that also has a *cis*-eQTL in at least one dataset on GeneNetwork, we performed a correlation analysis with all other probes or genes within that dataset. We then performed an enrichment analysis using WebGestalt using all the probes or genes that correlated with our gene of interest (i.e. the gene with a *cis*-eQTL), and investigated if any of the enriched annotations or pathways were relevant to the QTL.

Highly relevant enriched phenotypes were found in *9430012M22Rik* (location of *cis*-eQTL: Chr 3 55.291 Mb). This gene is present in QTL associated with vertical activity (BXD_12023). Genes that correlate with expression of *9430012M22Rik* in the neocortex are enriched for involvement in abnormal locomotor behavior (FDR=1.2056e-9) and abnormal voluntary movement (FDR=7.1848e-10). Other results for this gene that may be relevant include abnormal synaptic transmission and abnormal nervous system physiology [Supplementary table 7]. The genes that correlate with expression of *BC033915* (location of cis-eQTL: Chr9 45.671-48.081 Mb) in the hippocampus are enriched for abnormal motor capabilities/coordination/movement (FDR=2.3483E-11). Other relevant results include abnormal brain morphology and abnormal nervous system physiology. Similarly, this analysis has also revealed that genes expressing in the network of *Slitrk6* (location of *cis*-eQTL: Chr 14 109.231 Mb) in the striatum are involved in abnormal locomotor behavior (FDR=6.978E-12) and abnormal voluntary movement (2.9352E-11). Also, the phenotype of QTL containing *Slitrk6* is locomotion hence making this gene a good candidate.

Other genes with *cis*-eQTLs had significant enrinchments that include abnormal brain morphology, abnormal body composition and abnormal nervous system physiology [Supplementary table 7].

### Gene variant analysis

The second method that a variant can alter phenotype, is by changing protein structure or function. To examine this, we took advantage of the deep sequencing available for all BXD strains. We have identified over 5 million common SNPs and small INDELs which segregate within the BXD family (i.e. occur in greater than >20% of the population). For each of the 6 QTL identified above, we looked for variants that were predicted to alter protein structure or splicing, or predicted to be deleterious by SIFT or BLOSUM, using the variant effect predictor (VEP).

The Chr3:53.667-54.942Mb QTL for vertical activity contains predicted deleterious variants in 10 genes: two missense variants in *Ccdc169*; a missense variant in *Ccna1*; an inframe insertion in *Dclk1*; two frameshift variants, a stop loss, and nine missense variants in *Frem2*; a frameshift variant and six missense variants in *Mab21l1*; a frameshift variant, a missense variant, eight frameshift variants, two inframe deletions, 18 missense variants, and three stop losses in *Nbea*; four inframe deletions, an inframe insertion, six missense variants, and a start loss in *Postn*; a frameshift variant, eight missense variants, a stop gain, and a stop loss in *Spg20*; and a missense variant in *Trpc4*.

The Chr5:4.468-5.172Mb QTL for handling-induced convulsion in response to ethanol contains two missense variants in *Fzd1*, and four missense variants in *Cdk14*.

The Chr5:100.164-100.895Mb QTL for cocaine related phenotypes, contains predicted deleterious variants in 8 genes: three frameshift variants and three missense variants in *Cops4*; a missense variant and a stop-gain in *Enoph1*; a frameshift variant and five missense variants in *Hnrnpd*; three missense variants and a splice donor variant in *Hnrnpdl*; two frameshift variants, an inframe insertion, 13 missense variants, and a splice donor variant in *Hpse*; three missense variants in *LIN54*; a frameshift variant, one inframe deletions, and a splice donor variant in *Sec31a*; and a missense variant and a splice donor variant in *Tmem150c*.

The Chr9: 45.671-48.081Mb QTL for mechanical nociception contains predicted deleterious variants three genes: A frameshift variant, two missense variants and a stop loss in *4931429L15Rik*; two frameshift variants and five missense variants in *Cadm1*; and an inframe deletion, three missense variants, and a stop loss in *Cep164*.

The Chr11:62.923-65.082Mb QTL for nociception contains four frameshift variants, an inframe deletion, and thirteen missense variants in *Myocd*.

The Chr14:109.994-114.751Mb QTL for locomotion contains a stop loss, three frameshift variants, and 9 missense variants in *Slitrk6*.

### PheWAS analysis of the genes within QTLs

Another method to identify candidate genes, is to leverage data generated in another population, or another species. Phenome-wide association studies (PheWAS) take a gene or variant of interest, and find all reported associations in GWAS datasets. A number of these GWAS tools exist, using either different methods, or different human cohorts (https://atlas.ctglab.nl/PheWAS, http://pheweb.sph.umich.edu/).

Mouse QTL mapping has high power but low precision (i.e. we can detect a QTL, but do not know which of tens or hundreds of genes is causal), whereas human GWAS has low power but high precision (tens or hundreds of thousands of individuals are needed, but candidate regions are often smaller). By combining the power of mouse QTL mapping and the precision of human PheWAS, we are able to do more than either individually.

Candidate genes might show up in our analysis here that did not show up in our above analysis for several reasons, the most common being that gene expression was not measured in the relevant cell type or timepoint.

The QTL for vertical activity (Chr 3 51.723-56.473 Mb) includes a number of genes with relevant psychiatric, neurological and cognitive PheWAS hits. *Maml3* is associated with alcohol dependence [66] and depression [67]. *Cog6* has significant associations with Depressive symptoms [68] and worrier/anxious feelings [49]. *Nbea* is associated with nervous feelings [49] and alcohol dependence [66].

All the three genes present in the Chr5 4.468-5.172 Mb QTL (handling-induced convulsions, ethanol response) show significant PheWAS hits for psychiatric traits. *Fzd1* (location of cis-eQTL: Chr 5 4.753 Mb) is significantly associated with major depressive disorder [69].

In the Chr 5, 99.801-101.331 Mb region the genes *Hnrnpd* and *Lin54* show the highest number of relevant pheWAS hits. *Lin54* is associated with conditions like loneliness, anxiety, tension, and sleep related phenotypes [49,70,71].

*Cadm1* (Location of cis-eQTL: Chr 9 47.550 Mb) gene was found significantly associated with Schizophrenia and other psychiatric disorders [69,72].

Among the genes with *cis*-eQTLs in Chr 11, *Elac2* (location of *cis*-eQTL: Chr11 64.988 Mb) and *Arghap44* have most significant phenotype associations with schizophrenia/bipolar disorder [72,73].

The QTL for locomotion in the center (Chr14 109.994-114.751 Mb) shows two genes with PheWAS hits. *Slitrk6* is significantly associated with Parkinson’s Disease [74] and bipolar disorder [73]. *Slitrk5* has significant associations with various psychiatric traits including anxiety [49], nervous feelings [49] and alcohol dependence [66] [Supplementary Table 8].

## Discussion

We have demonstrated that old data in populations of isogenic strains can be reanalyzed, identifying novel genetic associations, containing novel candidate genes.

Of particular interest is Slitrk6 on Chr14. *Slitrk6* (SLIT And NTRK Like Family Member 6) is a protein coding gene. Our analysis strongly shows that abnormality in *Slitrk6* is implicated in disrupted locomotor behavior. The presence of *cis*-eQTL implies that a variant in this gene is effecting its expression and the gene is under its own regulation. Being part of a network in the striatum, whose genes are significantly involved in abnormal locomotory behavior and abnormal voluntary movement increases the plausibility. This gene is involved in altering both gene expression and protein structure/function. PheWAS analysis shows that this gene is involved in various neuropsychiatric and neurological phenotypes. The *Slitrks* have been previously mentioned as prominent candidate genes involved in neuropsychiatric disorders [75]. The members of the Slitrk family have been shown to be widely expressed in the central nervous system, with partially overlapping yet differential patterns of expression [76]. It is worth noting that this gene along with the other candidates have not been reported in Philip *et al*.’s study.

Another prominent finding is *Cadm1* (Cell adhesion molecule 1), a member of the immunoglobulin superfamily, present on Chr9. Its shows the presence of a *cis*-eQTL and is found to be associated with schizophrenia. *Cadm1* knockout mice show anxiety-like behavior under certain conditions like the open-field and light-dark transition tests and motor coordination and gait were impaired in rotarod and footprint tests [61]. The role of CADM1 in relation to prefrontal brain activities, inhibition function, and ADHD, indicating a potential “gene–brain–behavior” relationship was shown by a study that evaluated the association of CADM1 genotype with ADHD, executive function, and regional brain functions [77]. Studies show a connection between ADHD and pain tolerance [78]. Adults with ADHD are comparatively more sensitive to pain. In such cases, Dopamine agonists like methylphenidate (MP) may exert antinociceptive properties [79] and normalize pain perception. Adults and children with ADHD exhibit motor regulation problems which are in turn associated with the pain levels [80].

We discovered a novel QTL regulating handling induced convulsions after ethanol treatment (BXD_11635) on Chr5:4.468-5.172. Only three genes are within the confidence interval for this QTL, and two of which, *Fzd1* and *Cdk14*, have *cis*-eQTL and predicted deleterious variants. Interestingly, *Cdk14* is regulated by the anticonvulsant drug valproic acid [81,82,83], and is up-regulated in malaria patients who experience febrile convulsions [84].

We have shown that by using families of isogenic strains, we can not only go back and discover new phenotype-genotype associations that were not previously found, but that we can find highly plausible candidate genes within these novel QTL.

## Supplementary Materials

Supplementary table 1: Phenotype data and reanalysis with new tools and genotypes

For each of the traits obtained from Philip et al. 2010, H-K mapping results from Philip’s study and corresponding GEMMA results of Max −log(p), Max −log(p) Peak location and Max LRS value from our study are shown.

Supplementary table 2: Significant traits with QTL regions.

QTL confidence intervals (QTL start region and QTL end region) for each of significant traits determined by 1.5 LOD or 1.5 −log(p) drop and number of genes in each of these regions are shown.

Supplementary table 3: Significant traits (H-K Vs GEMMA and R/qtl2)

A total of 426 significant traits which had a maximum LRS < 17 with H-K (i.e. non-significant), and now have a maximum −log(p) > 4 (significant with GEMMA). 61 traits of these are significant with R/qtl2 with permutations, as mentioned in the table.

Supplementary table 4: QTLs and respective summaries

QTLs were segregated and identified from significant, independent traits. 25 Novel QTLs detected in this study with the number of genes and phenotypes of these QTLs are shown.

Supplementary table 5: Comparison of QTLs found in this study with Philip et al. and other studies.

The novel QTL found in our study compared to the relevant abnormal behavioral phenotypes found in those QTLs in Philip’s study and other studies, if any, are shown.

Supplementary table 6: Genes and cis-eQTLs

Six small QTLs with less than a 100 genes considered for cis-eQTL analysis. Each gene in these QTLs analysed for the presence of cis-eQTL are shown with the location. Correlation matrix generated between the genes in a QTL with cis-eQTL in any brain tissue and the phenotype of that QTL. Brain tissues with highest correlation are also shown.

Supplementary table 7: Coexpression networks or ‘gene-friends’

The correlations of genes with cis-eQTLs in the regions of highest correlated brain tissue are analysed in Webgestalt for enriched annotations and pathways. The enrichment results, FDR value, P-value and enrichment ratio are shown for each gene with cis-eQTL.

Supplementary table 8: PheWAS

Results of PheWAS analysis of all the genes present in the small QTLs of our interest are shown containing the psychiatric, neurological and cognitive traits that may be the most relevant to our study.

---

**Table 1:**
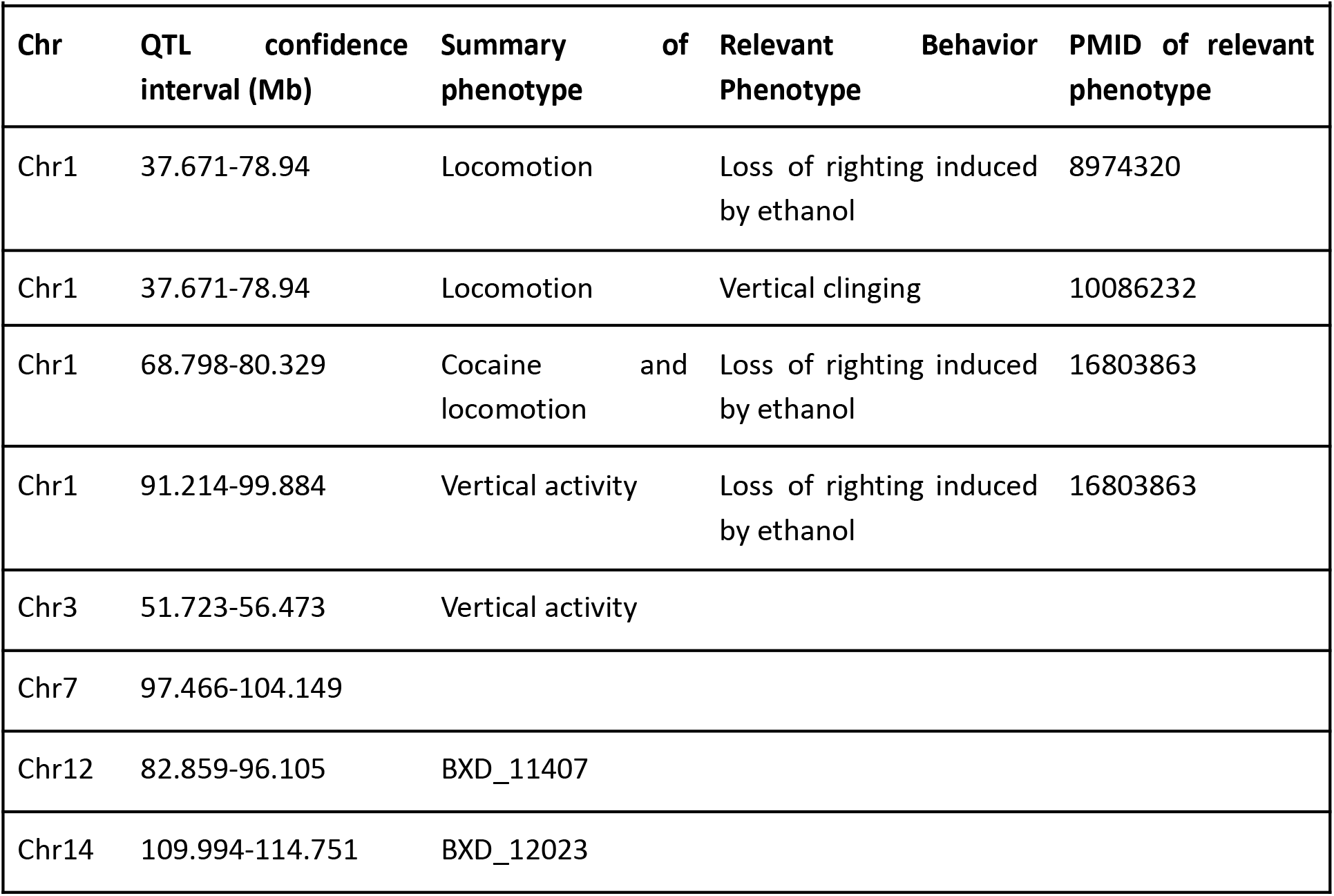
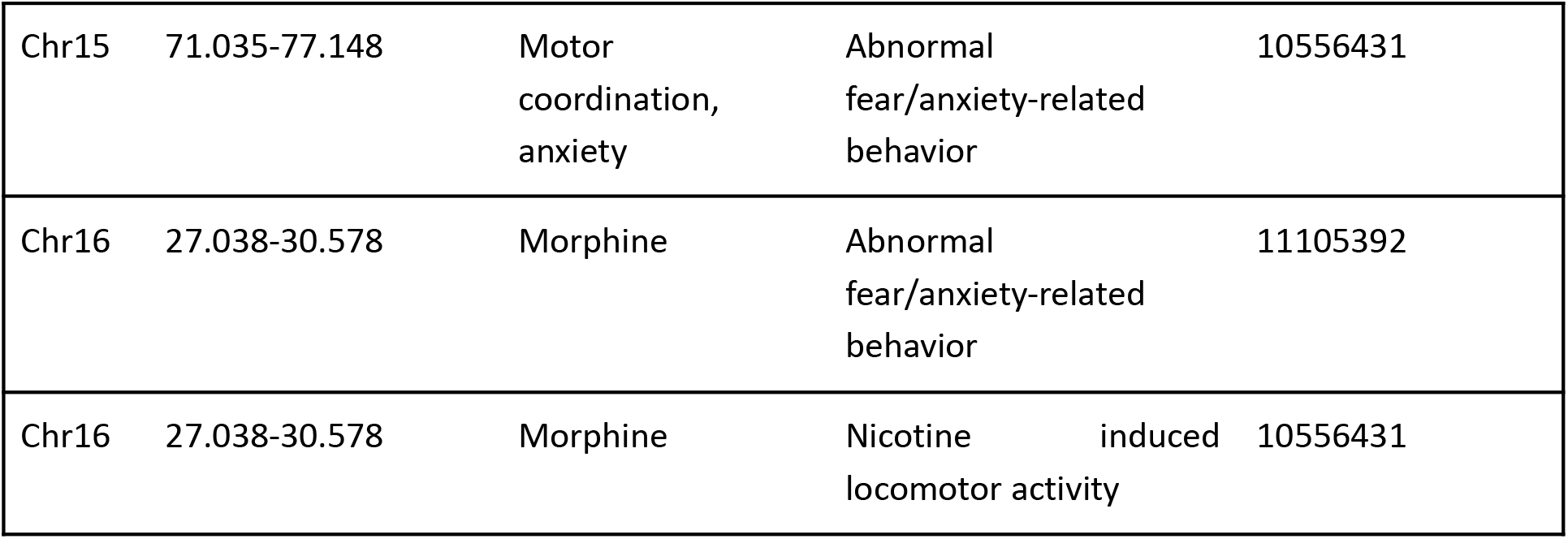
Summary of novel QTL, not found at the significant or suggestive level in the original paper by Philip et al [39]. The position of the QTL, a summary of the phenotypes within that QTL, and relevant phenotypes found in other studies are shown. Details of all identified QTL are in Supplementary table 5.

